# A high-throughput segregation analysis identifies the sex chromosomes of *Cannabis sativa*

**DOI:** 10.1101/721324

**Authors:** Djivan Prentout, Olga Razumova, Bénédicte Rhoné, Hélène Badouin, Hélène Henri, Cong Feng, Jos Käfer, Gennady Karlov, Gabriel AB Marais

**Author notes:** corresponding author: Gabriel Marais; LBBE – UMR 5558, CNRS / Université Lyon 1, campus de la doua, 69622 Villeurbanne cedex, France.

## Abstract

*Cannabis sativa*-derived tetrahydrocannabinol (THC) production is increasing very fast worldwide. *C. sativa* is a dioecious plant with XY chromosomes, and only females (XX) are useful for THC production. The *C. sativa* sex chromosomes sequence would improve early sexing and better management of this crop; however, the *C. sativa* genome projects failed to identify the sex chromosomes so far. Moreover, dioecy in the Cannabaceae family is ancestral, *C. sativa* sex chromosomes are potentially old and thus very interesting to study as little is known about the last steps of sex chromosome evolution in plants. Here we RNA-sequenced a *C. sativa* family (2 parents and 10 male and female offspring) and performed a segregation analysis for all *C. sativa* genes using the probabilistic method SEX-DETector. We identified >500 sex-linked genes. Mapping of these sex-linked genes to a *C. sativa* genome assembly identified a single chromosome pair with a large non-recombining region. Further analysis of the >500 sex-linked genes revealed that *C. sativa* has a strongly degenerated Y chromosome and represents the oldest plant sex chromosome system documented so far. Our study revealed that old plant sex chromosomes can have large non-recombining regions and be very differentiated and still be of similar size (homomorphic).

## INTRODUCTION

*Cannabis sativa* is an ancient crop (Schultes et al. 1974) with two main traditional uses: marijuana and hemp (Small 2015). Marijuana, which is used in folk medicine, as a recreational drug, and lately in conventional medicine (Alexander 2016), has a narcotic effect thanks to tetrahydrocannabinol (THC) and other cannabinoids produced in high concentration by some *C. sativa* cultivars. Until recently, the use of marijuana was prohibited in almost all countries, but *C. sativa*-derived products with high THC concentrations are now legal in several US states, Australia, Germany, Peru and the UK for medicinal purposes (Offord 2018) and also in Uruguay, Canada and several US states for recreational use (Yeager 2018). In the US, marijuana legal economy amounted ∼$17 billion in 2016 and may reach as much as $70 billion / year by 2021 (McVey 2017). However, legalisation of marijuana is so recent that very few biotech tools have been developed for high THC-producing *C. sativa* cultivars (Yeager 2018). Interest on hemp is also increasingly attractive as crop for the sustainable production of fibres and oils (Andre et al. 2016; Salentijn et al. 2019). Hemp cultivars usually have a low level of THC and can legally grow in many countries where marijuana is illegal. Features of male and female hemp plants differ and early sexing is also wanted.

THC reaches the highest concentrations in female inflorescences (bracts), so that only female *C. sativa* plants are of economical importance; furthermore, pollinated female plants produce smaller inflorescences and therefore less THC (Small 2015). It is thus important to avoid growing male plants as they are a waste of resources, labour and space. Sexual dimorphism in *C. sativa* is weak as in many dioecious plants (Barrett and Hough 2013), and sex can be determined with certainty only when the plants start flowering (Small 2015). *C. sativa* is a dioecious plant in which sex is determined by a XY chromosome system (Divashuk et al. 2014). Indeed, a few Y-linked genetic markers have been identified in the past and are used to sex *C. sativa* seedlings (e.g (Techen et al. 2010)). However, it is not known whether these markers work with all cultivars. The *C. sativa* sex chromosomes sequences would thus be an important genomic resource that could help improving agricultural yields. However, the *C. sativa* genome projects (van Bakel et al. 2011; Grassa et al. 2018; Laverty et al. 2019) have failed to identify the sex chromosomes, despite chromosome-level assemblies in the latest projects.

*C. sativa* is one of 15,600 dioecious species of flowering plants (Renner 2014). Dioecy and sex chromosomes have evolved multiple times in plants (Renner 2014) but very few plant systems have been studied in details (Ming et al. 2011; Charlesworth 2015; Muyle et al. 2017) and the current model for the evolution of plant sex chromosomes includes many gaps, the late stages in particular have not been studied. Old animal sex chromosomes systems are usually heteromorphic with the Y being smaller than the X (Charlesworth et al. 2005; Bachtrog 2013). Heteromorphic systems in plants are much more recent and in the case of *Silene latifolia* and *Coccinia grandis*, the Y is larger than the X (Matsunaga et al. 1994; Sousa et al. 2013), probably due to accumulation of DNA repeats on the Y (Sousa et al. 2016; Hobza et al. 2017). The Cannabaceae and related families (Urticaceae, Moraceae) family is expected to derive from a dioecious common ancestor (Zhang et al. 2019). The sex chromosomes of *C. sativa* could thus be much older than those of the species studied so far. Interestingly, *C. sativa* X and Y chromosomes are of similar size (= homomorphic, see (Divashuk et al. 2014)).

Here we used a recently developed statistical tool to identify X- and Y-linked genes, called SEX-DETector (Muyle et al. 2016). SEX-DETector analyses genotyping data from a cross (2 parents and a few offspring individuals, see Figure 1). Patterns of allele transmission from parents to progeny differ for an autosomal or a sex-linked gene. For example, an allele only transmitted from father to sons is clearly indicative of a Y-linked allele. SEX-DETEctor relies on a probabilistic model that accounts for typical errors in genotyping data and is used to compute, for each gene, the probability of autosomal and sex-linked segregation types. This key feature of SEX-DETEctor makes it better at making inferences about segregation type than an empirical approach relying on data filtering to remove genotyping errors would do (better sensitivity, similar specificity, see (Muyle et al. 2016)). We applied SEX-DETector to *C. sativa*, inferred sex-linked genes and used those genes to 1) identify the sex chromosomes of *C. sativa* in an available reference genome assembly 2) characterize the *C. sativa* XY system and compare it to other plant systems.

**Figure 1:**
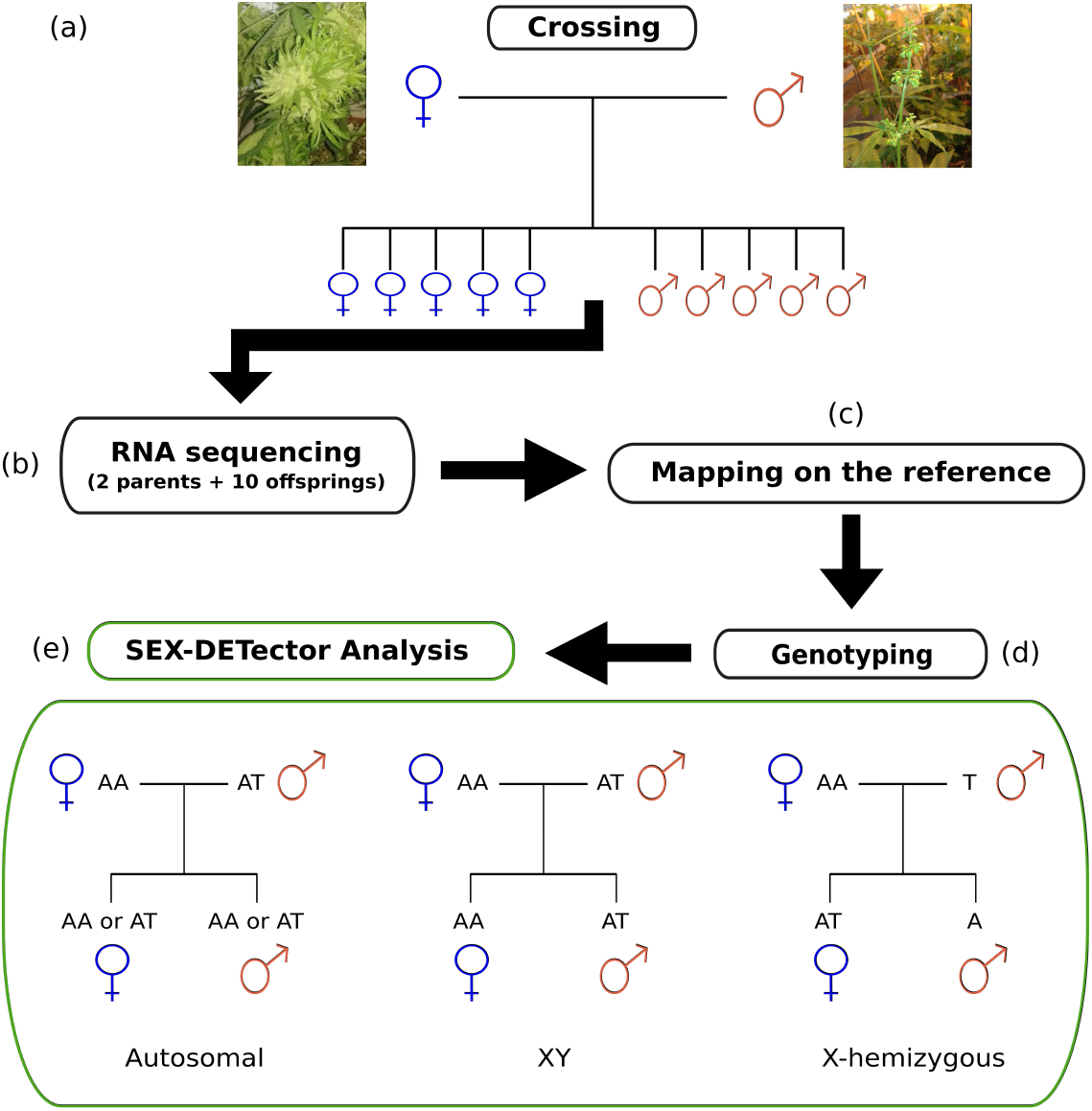
Experimental design and bioinformatic pipeline to identify sex-linked genes. The SEX-DETector analysis relies on obtaining genotyping data from a cross (parents + F1 progeny). It infers the segregation type based on how alleles are transmitted from parents to offspring. Examples of segregation patterns are shown here for the three segregation types included in SEX-DETector. See Methods for more information.

## RESULTS

### Identifying sex-linked genes in *C. sativa*

More than 576 millions 50 bp single-end reads of the parents and 5 male and 5 female offspring were mapped on the reference transcriptome of (van Bakel et al. 2011), and all individuals were genotyped (see Methods). Of these, 11.515 were inferred as autosomal, and 565 genes were identified as sex-linked. The latter included 347 X/Y gene pairs and 218 X-hemizygous genes (*i.e.* genes lacking Y copies, see Methods and Table 1). The sex-linked genes represented 4.6% of the genes for which SEX-DETector produced an assignment.

**Table 1:** Results of the SEX-Detector analysis.

|  | Numbers |
| --- | --- |
| All genes* | 30,074 |
| Genes with at least 1 SNP detected, used for SEX-DETECTOR analysis | 28,456 |
| Genes with undetermined segregation type** | 16,381 |
| Autosomal genes | 11,510 |
| All sex-linked genes | 565 |
| X/Y gene pairs | 347 |
| X-hemizygous genes | 218 |
| Estimated Y gene loss rate | 70% |

\* transcripts from the gene annotation of the reference genome, see Methods
\*\* lack of information

### Identifying the sex chromosome pair in *C. sativa*

A total of 363 sex-linked genes (out of the 555 that we could map) mapped to the chromosome 1 in the reference genome (XY gene pairs: 166/340 = 48.8%, X-hemizygous genes: 197/215 = 91.6%, Figure 2), which points to chromosome pair number 1 being the sex chromosome pair. 192 sex-linked genes (i.e 36% of all sex-linked genes) mapped to other chromosomes (discussed below). Sex chromosomes typically have non-recombining regions in which the synonymous divergence between the X and Y copies of a sex-linked gene (also called gametologs) can be substantial (Charlesworth 2015; Muyle et al. 2017).

**Figure 2:**
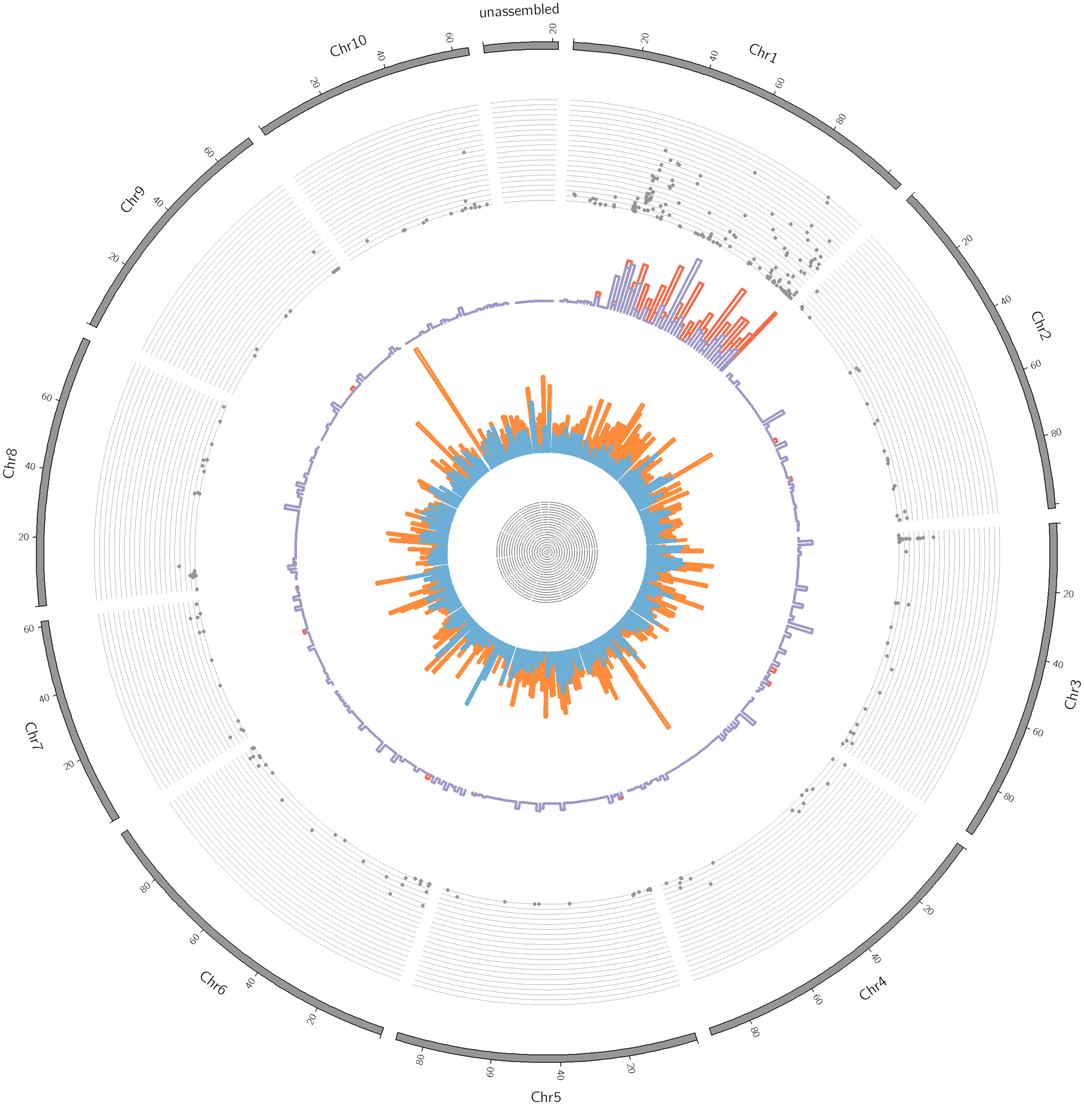
Distribution of the sex-linked and sex-biased genes onto the *C. sativa* reference genome. From outer to inner rings: 1) chromosomes from 1 to 10 and unassembled scaffolds of the reference genome (Grassa et al. 2018), 2) X-Y dS values (from 0 to 0.4), 3) cumulative distribution of the XY-linked genes (in blue) and X-hemizygous genes (in red) corrected for gene density (proportion of sex-linked genes among all genes)), 4) distribution of genes with sex-biased expression: male-biased (light blue), female-biased (orange) (proportion of sex-biased genes among all genes).

Using the sex-linked SNPs inferred by SEX-DETector, we are able to quantify the synonymous divergence (dS) between X and Y copies, which reaches 0.4 in the mostly divergent XY gene pairs (Figure 3A). We found that the most divergent XY gene pairs (showing the highest X-Y dS values) mapped to the chromosome 1 as expected if this chromosome pair is the sex chromosome pair. We also found that two regions of the chromosome 1 could be distinguished (Figure 2): region 1 (from 30 to 105 Mb) where the XY gene pairs with the highest dS values are found (mean X-Y dS = 0.079, top 5% X-Y dS = 0.32, top 10% X-Y dS = 0.28), and where 58.6% of the sex-linked genes of the region are X-hemizygous (*i.e* having potentially lost their Y copies), and region 2 (from 1 to 30 Mb) showing little divergence (mean X-Y dS = 0.014, top 5% X-Y dS = 0.05, top 10% X-Y dS = 0.04) and 9.3 % are X-hemizygous genes. These observations suggest region 1 is the X-specific region (not recombining in males), and region 2 the pseudo-autosomal (still recombining in males) one.

**Figure 3:**
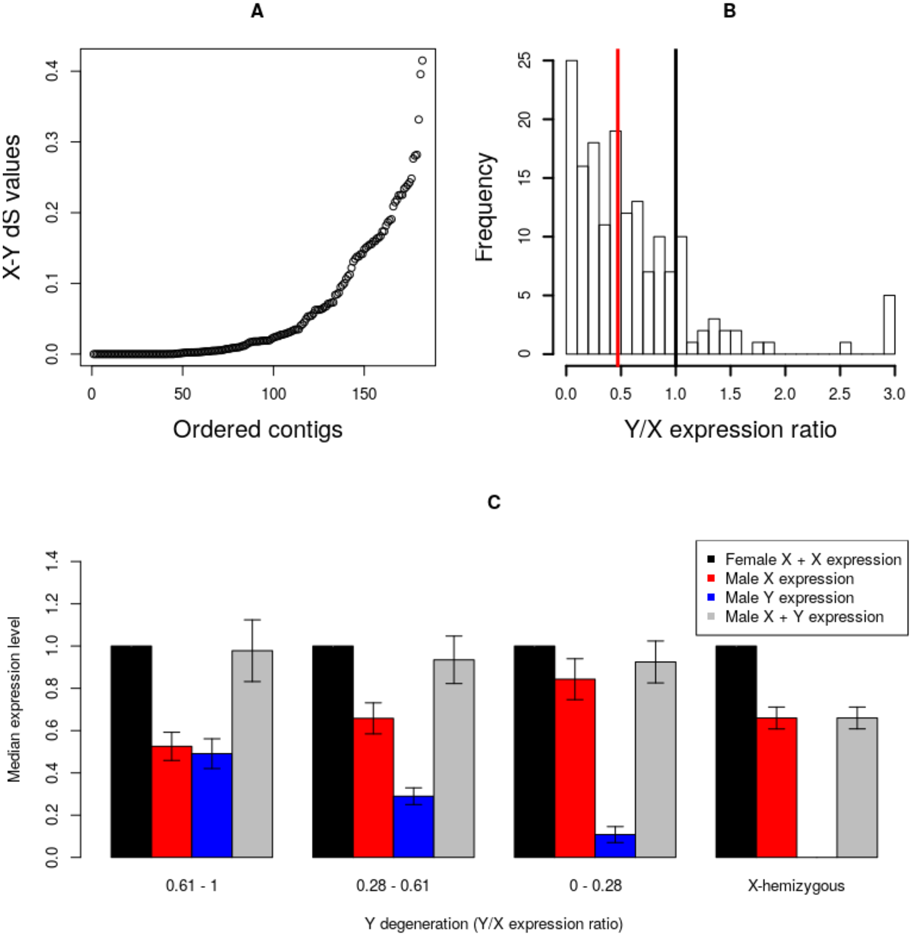
Patterns of molecular evolution of *C. sativa* sex chromosomes. A) X-Y dS values for the XY gene pairs, B) Y/X expression ratio for the XY gene pairs, the black bar shows the expected value for no Y degeneration, the red bar shows the median observed here (median = 0.47). Both are significantly different (Wilcoxon paired test p-value = 2.2 × 10^-16^). C) Dosage compensation in *C. sativa*. The expression levels of the X and Y alleles in males and females are shown for gene categories (from left to right, categories 0.61-1 and 0.28-0.61: N = 44, category 0-0.28: N = 43, X-hemizygous: N = 218) with different levels of Y degeneration (measured by the Y/X expression ratio). Sex-biased genes (with strong and significant differences in male and female expression) have been removed, as they are not expected to exhibit dosage compensation (see (Muyle et al. 2012; Muyle et al. 2018)). Only sex-linked genes mapping to chromosome 1 have been included here. Figure S1 shows the same analyses with all sex-linked genes.

### Age of the *C. sativa* sex chromosome system

We then used the 565 sex-linked genes to study the evolution of sex chromosomes in *C. sativa*. First, we used the dS values of the X/Y gametologs and different molecular clock estimates for plants to infer the age of the sex chromosomes on *C. sativa* (see Methods). Using the maximum observed dS value (0.4), we estimated that recombination suppression between X and Y chromosomes have initiated 26.7-28.6 My ago in *C. sativa*. If we use the dS values of the 5% or 10% most divergent gene pairs to be more conservative when estimating the maximum X-Y divergence, we obtain more recent ages for the initial recombination suppression (17.3-18.6 My old using top 5% X-Y; 12-13 My old using top 10% X-Y, see Table 2).

**Table 2:**
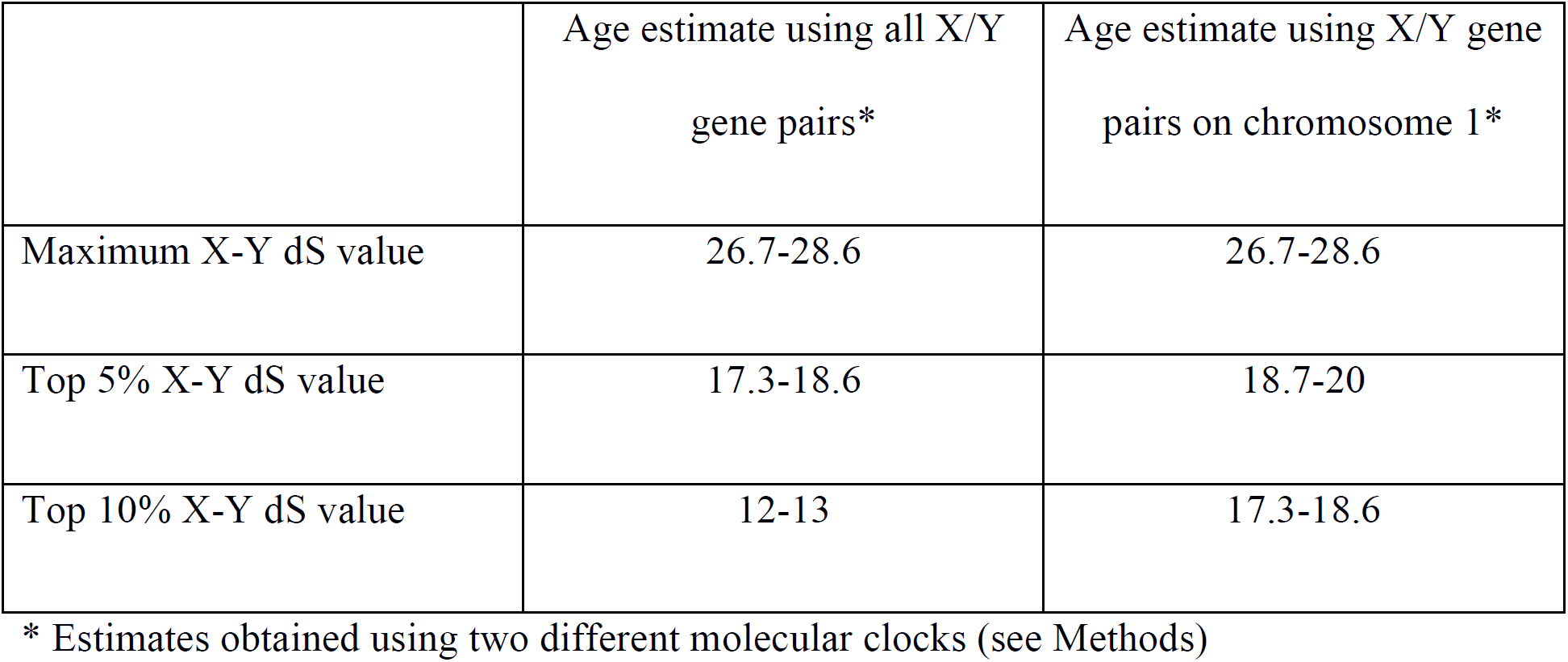
Estimates of the age of the *C. sativa* sex chromosome system.

### Degeneration of the Y chromosome and dosage compensation in *C. sativa*

Second, we studied the extent of Y degeneration in *C. sativa* and estimated gene loss using the X-hemizygous genes (see Methods). This measure of Y gene loss is of course a rough estimate as it reflects both true loss and simply the absence of expression of the Y copy in flower buds (Bergero and Charlesworth 2011). The number of X-hemizygous genes is known to be underestimated with respect to that of XY gene pairs, as X-hemizygous genes can only be detected when there is polymorphism in the X. To correct for this, we compared the number of X-hemizygous genes (218) and the XY gene pairs with polymorphism in the X copy (89), and we found that ∼70% of the Y-linked genes may have been lost in *C. sativa*. The results were similar when focusing on sex-linked genes found on chromosome 1 only (72.5%).

To study further Y degeneration, we focused on the expression of the sex-linked genes. Allele-specific expression analysis at the X/Y gene pairs revealed a median Y/X expression ratio of 0.50 overall (347 genes) and 0.47 for chromosome 1 genes only (166 genes, see Figure 3B), much lower than the expected 1.0 value in case of equal Y/X expression (*i.e* no Y degeneration). We found some evidence for dosage compensation as, in males, expression of X was increased when expression of Y was reduced (Figure 3C). The results were unchanged when using all inferred sex-linked genes or only those found on chromosome 1 (Figure 3 and S3).

### Genomic distribution of the sex-biased genes in *C. sativa*

15.7% of the genes expressed in flower buds are differentially expressed between male and female individuals (sex-biased genes, see Table 3 and Figure S2). The male-biased genes are significantly more numerous than the female-biased genes (9.06% vs. 6.64%, Fisher exact test p-value = 2.2 × 10^-16^, see Table 3), and sex-biased genes are distributed all over the *C. sativa* genome (see Figure 2). The sex-linked genes were significantly enriched among the sex-biased genes (25.8%) compared with the autosomal genes (13.9%; Fisher exact test p-value = 3.7 × 10^-13^).

**Table 3:** List of sex-biased genes.

| Gene categories | Total | Female-biased expression | Male-biased expression | p-value* |
| --- | --- | --- | --- | --- |
| All genes with sex biased-expression | 3483 | 1473 | 2010 | $2.2 \times 10^{-16}$ |
| Autosomal genes | 1599 | 725 | 874 | $2.1 \times 10^{-4}$ |
| Sex-linked genes | 146 | 79 | 67 | 0.36 |
| XY gene pairs | 87 | 34 | 53 | 0.053 |
| X-hemizygous genes | 59 | 45 | 14 | $6.5 \times 10^{-05}$ |

\* Exact binomial test, with a theoretical mean equal to 0.5

## DISCUSSION

### Chromosome pair 1 is the sex chromosome pair in *C. sativa*

Using SEX-DETector, we have been able to identify a large number of sex-linked genes with a moderate sequencing effort (576 millions of single-end 50 bp reads). We identified a chromosome pair (chromosome 1 in the assembly of (Grassa et al. 2018)) as the sex chromosomes of *C. sativa*. A part of the sex-linked genes did not map to the chromosome 1, they could be false positives or result from assembly errors (see Text S1).

### The sex chromosomes are the largest in the *C. sativa* genome

This pair is the largest pair of the *C. sativa* genome, in agreement with cytogenetic data (Divashuk et al. 2014). This is frequent in plants with heteromorphic sex chromosomes (X and Y have different size, e.g. *S. latifolia, C. grandis*, see (Muyle et al. 2017) for review), but also in species with homomorphic chromosomes (X and Y have similar size) such as papaya, where both the X and Y increased in size due to the accumulation of repeats (Gschwend et al. 2012; Wang et al. 2012). We did not observe signs of such a process in *C. sativa*, as the gene density on the X chromosome is similar to the gene density on the autosomes (32 genes / Mb vs. 33 genes / Mb). It is thus possible that the sex chromosomes are the largest in *C. sativa* simply because the sex-determining genes happened to evolve on the largest pair of chromosomes.

### *C. sativa* sex chromosome are old

Age estimates of the *C. sativa* sex chromosome ranges from ∼12 My to ∼29 My old (Table 2). This is possible given that dioecy probably is ancestral for the whole Cannabaceae family that diversified ∼80 My ago (Zhang et al. 2019). They may be the oldest sex chromosomes in plants for which the age was inferred from sequence data (Ming et al. 2011; Charlesworth 2015; Muyle et al. 2017). For instance, sex chromosomes are ∼11 My old in *S. latifolia* (Krasovec et al. 2018) and 8-16 My old in two dioecious *Rumex* species (Crowson et al. 2017).

Further evidence that the *C. sativa* sex chromosomes are older than those of *S. latifolia* and *R. hastatulus* is the fact that the median Y/X expression ratio is ∼0.5, much lower than what has been reported for the other species (∼0.8 for *S. latifolia* (Bergero and Charlesworth 2011; Muyle et al. 2012) and ∼0.8 for the old sex-linked genes *R. hastatulus* (Hough et al. 2014)), and that gene loss is about 70%, which is very higher than those species (∼40% for *S. latifolia* (Papadopulos et al. 2015; Muyle et al. 2018) and 30% in *R. hastatulus* (Hough et al. 2014)). In *R. rothschildianus*, gene loss amounts ∼90% but the degeneration speed not the age of the system is believed to explain this observation (Crowson et al. 2017)). Thus, the Y chromosome of *C. sativa* seems more strongly degenerated than the Y chromosomes of species with strong heteromorphism.

### Implications for the model for sex chromosome evolution in plants

Most of the plant sex chromosome systems that have been studied so far either have small non-recombining regions and homomorphic sex chromosomes (e.g. *Carica papaya, Asparagus officinalis, Diospyros lotus*), or have heteromorphic sex chromosomes with the Y being larger than the X (e.g. *Silene latifolia, Coccinia grandis*). We here found that in a species with (nearly) homomorphic sex chromosomes, the non-recombining region is large, as it represents ∼70% (85 / 105 Mb) of the *C. sativa* sex chromosomes (as suggested in (Divashuk et al. 2014) based on cytogenetic data).

In the current scenario for the evolution of the sex chromosomes in plants (Ming et al. 2011; Charlesworth 2015; Muyle et al. 2017), it is unclear where these XY chromosomes fit. Indeed, sex chromosome evolution in plants is thought to start with a small non-recombining region on the Y chromosome, which accumulates DNA repeats and tends to grow (Figure 4). In papaya, the Y non-recombining region is ∼8 Mb large while the X homologous region is ∼4 Mb (Wang et al. 2012). In some dioecious plants, DNA repeat accumulation in the Y non-recombining region has been fast and Y chromosomes that are much larger than the X have evolved in *Silene latifolia* (Matsunaga et al. 1994) and *Coccinia grandis* (Sousa et al. 2013).

**Figure 4:**
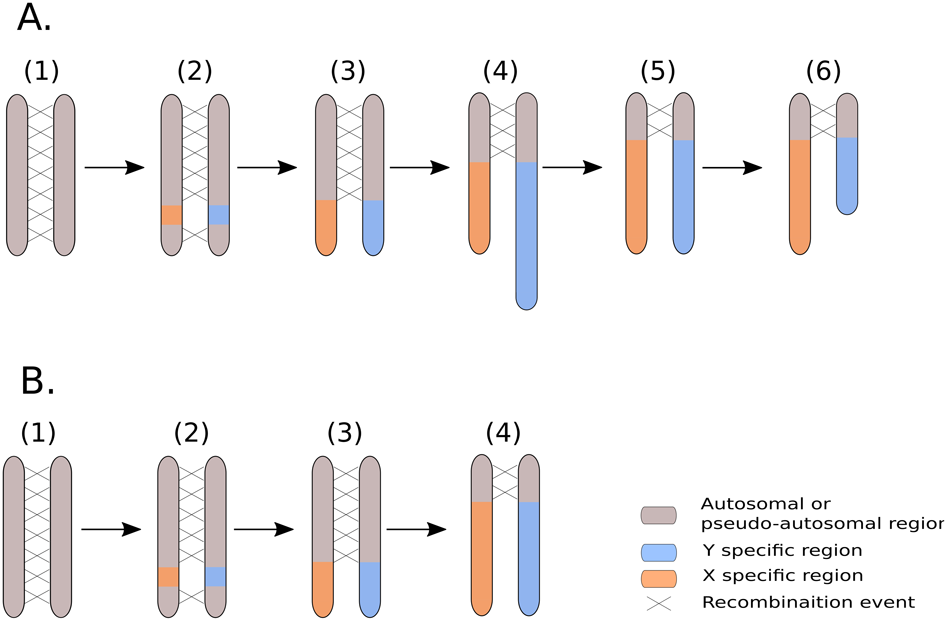
Revisiting the model for the evolution of plant sex chromosomes with *C. sativa*. A) The current model for the evolution of plant sex chromosomes is as follows: 1) Sex chromosomes originate from autosomes on which sex-determining genes evolve, 2) The region encompassing the sex-determining genes stops recombining, 3) the non-recombining region grows larger due to additional events of recombination suppression, 4) the non-recombining region of the Y chromosome accumulates repeats and can become larger than the corresponding region on the X chromosome, 5-6) the Y chromosome undergoes large deletions, and ultimately becomes smaller than the X chromosome. Steps 1-4 have been previously documented in plants (e.g. (Charlesworth et al. 2005; Ming et al. 2011; Muyle et al. 2017) for review) while steps 5-6 are very speculative. Our study is supportive of this scenario if we assume that the *C. sativa* Y chromosome has been larger in the past. B) It is possible however that the accumulation of repeats has been slow in the Y chromosome of the *C. sativa* lineage and that X and Y chromosomes have always been of similar size (see text). Here step 4 does not imply the elongation of Y chromosome.

Either DNA repeat accumulation on the Y has been slow in the *C. sativa* lineage, or the Y used to be larger than it is today and has undergone genomic shrinking, a process that is reminiscent of the evolution of the sex chromosomes in animals (Ming et al. 2011; Bachtrog 2013), where old Y chromosomes can be tiny compared to their X counterpart (Figure 4). Distinct assemblies for the X and Y chromosomes in *C. sativa* and also sequencing of other dioecious Cannabaceae species will help testing this idea in the future.

## METHODS

### Plant material, RNA extraction, and sequencing

One male and one female *C. sativa* plants (‘Zenista’ cultivar) were grown in controlled conditions in greenhouse. A female was crossed with a male plant (controlled pollination). Seeds from this cross were sown to produce the F1. Flower buds were sampled from the parents and 5 offspring of each sex as in (Muyle et al. 2012). Total RNA was isolated from young floral buds using the “RNeasy Plant Mini Kit” (Qiagen) plant isolation kit as recommended by the producer. After isolation of the RNA was placed in tubes RNAstable® (Sigma). One library per individual was prepared. RNA-sequencing was conducted using the Complete Genomic (CG) technology, which provides 20 millions ∼50 bp single-end reads per sample (Liu et al. 2012). Two CG runs were done and we obtained a mean of 48 millions reads per individuals (see Table S1). Read quality was good (phred score > 35 for all reads) and no trimming was performed.

### Mapping, genotyping and SEX-DETector analysis

The SEX-DETector analysis requires mapping the reads of the individuals onto a reference transcriptome and performing a SNP-calling to genotype all individuals for all expressed genes. We extracted the 30,074 transcripts from the annotation of the 2011 complete genome (van Bakel et al. 2011). The initial mapping analyses were done using BWA, allowing for 5 mismatches per read (version 0.7.15-r1140, bwa aln -n 5 and bwa samse, see (Li and Durbin 2010)) and Bowtie2 (version 2.1.0, bowtie2-build and bowtie2 –x, see (Langmead and Salzberg 2012)). We used Samtools (version 1.3.1, samtools view -t output.fa -F 4 -h and samtools sort -m 2G, see (Li et al. 2009)) to convert files from the mapping .bam files into.sam files and to remove unmapped reads.

The genotyping was performed using reads2snp (version 2.0.64, reads2snp -aeb -min 3 -par 0, see (Gayral et al. 2013)), as recommended (Muyle et al. 2016) (i.e. by accounting for allelic expression biases and without filtering for paralogous SNPs). Only SNPs supported by at least three reads were conserved for subsequent analysis (except in Table S2).

We ran SEX-DETector (-system xy/zw/no_sex_chr -seq -detail -detail-sex-linked -L -SEM - thr 0.8, see (Muyle et al. 2016)) on genotyping data of the 12 individuals. SEX-DETector uses a maximum likelihood approach to estimate the parameters of its model, which include several genotyping error parameters. The posterior probability of being autosomal (P_A), XY (P_XY) or X-hemizygous (P_Xh) is then computed for each SNP and then computed for each transcript (combining the posterior probabilities of all SNPs, see (Muyle et al. 2016)). A transcript was inferred as sex-linked if its posterior probability of being either XY or X-hemizygous was ≥ 0.8 (i.e. P_XY + P_Xh ≥ 0.8) and if at least one sex-linked SNP had no genotyping error; autosomal segregation was inferred similarly (P_A ≥ 0.8 and at least one autosomal SNP without genotyping error; see (Muyle et al. 2016)). The remaining transcripts were considered undetermined and were not used for further analysis. Among the sex-linked transcripts, those with an expressed Y-allele were considered XY, the others X-hemizygous.

SEX-DETector runs on the first mapping with BWA and Bowtie2 yielded high Y genotyping error (YGE) parameter values, which could be the result of mapping errors of Y-linked reads (Muyle et al. 2016). The reference transcriptome used for mapping was derived from the genome of a female plant (van Bakel et al. 2011), which may result in a mapping bias against the Y-linked reads. To solve this problem, we used GSNAP (version 2017-11-15, gsnap -m 5, see (Wu and Nacu 2010)), which can be used to map RNA-seq reads onto a divergent reference. GSNAP was thus used in a SNP-informed mode that adjusts read alignment onto a reference taking into account a user-provided list of SNPs that are not considered as mismatches. For this procedure, we first mapped reads with BWA and collected all the SNPs present in SEX-DETector output, which were provided to GSNAP. We ran four iterations of GSNAP. For each iteration, SEX-DETector detected new sex-linked SNPs, which were added to the list of SNPs provided to GSNAP. As expected, the Y genotyping error parameter value decreased from 0.84 with BWA to 0.07 with 4th GSNAP iteration (Table S2), and the mapping rate from 82.57% to 87% (Table S1).

### Circular representations of location of sex-linked genes in the *C. sativa* genome

To map the sex-linked genes onto the *C. sativa* genome, we used blast to find the best hit of each *C. sativa* transcript in the 2011 transcriptome on a recent (unannotated) reference genome (blastn -max_target_seqs 1 -max_hsps 1). For this mapping, we used the *C. sativa* reference genome with the best assembly statistics (size = 875 Mb, 10 pseudomolecules, 220 scaffolds, N50 = 91 Mb, see https://www.ncbi.nlm.nih.gov/genome/genomes/11681 and (Grassa et al. 2018)). We then used Circos (version 0.69-6) for visualizing the location of sex-linked genes. The total gene number per window along the *C. sativa* genome was obtained by dividing the number of mapped transcripts by window size. Bedtools makewindows (version v2.26.0) was used for the separation of the genome in windows of 2Mb and bedtools intersect (version v2.26.0, -c option) for computing the gene number per window.

### Analysis of the sex-linked genes

#### Y gene loss

To estimate the rate of gene loss in the Y chromosome, we compared the number of X/Y gene pairs and the number of X-hemizygous genes as in (Bergero and Charlesworth 2011). Identifying X/Y gene pairs relies on fixed X/Y differences while identifying X-hemizygous genes relies on X-polymorphism only, which makes detection of X-hemizygous genes less likely (see (Bergero and Charlesworth 2011; Muyle et al. 2016)). Y gene loss proportion estimate was thus corrected for this bias as follows:

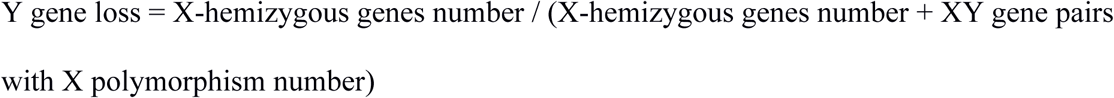

#### Values of synonymous divergence (dS) and age of the XY system

The X and Y open reading frames sequences were aligned using the translated reference transcripts to get reading-frame informed alignments. X-Y dS values were obtained using codeml (PAML version 4.9, see (Yang 2007)) in pairwise mode. To estimate the age of the *C. sativa* XY system, we considered maximum X-Y dS values and used two different molecular clocks for plants: 1.5 × 10^-8^ substitutions / site / year (Koch et al. 2000) and 7 × 10^-9^ mutations / site / generation (Ossowski et al. 2010). We obtained age of the XY system as follows:

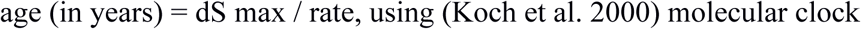

age (in number of generations) = dS max / 2 µ, using (Ossowski et al. 2010) molecular clock; the age in million years was obtained assuming 1 generation per year in natural populations of *C. sativa*.

#### Allele-specific expression analysis

We used allele-specific expression estimates at X/Y gene pairs provided by SEX-DETector (Muyle et al. 2016) for the estimation of Y/X expression ratio and patterns of dosage compensation (see Figure 3B-C). These estimates relied on counting reads spanning X/Y SNPs only and were normalized using the total read number in a library for each individual. These estimates were further normalized by the median autosomal expression for each individual.

### Identifying sex-biased genes

As the differential gene expression analysis methods currently available varied in performance (Schurch et al. 2016; Costa-Silva et al. 2017), we chose to combine several methods. Analysis contrasting the gene expression level between our 12 male and female individuals were thus performed using three R packages (i) DEseq2 version 1.10.1 (Love et al. 2014), (ii) EdgeR version 3.26.9 (Robinson et al. 2010) both relying on negative binomial distribution of read count modelling and (iii) limma-voom version 3.26.9 (Ritchie et al. 2015) based on log-normal distribution modelling to take into account the sampling variance of small read counts. Very low expressed genes were discarded from the analysis, keeping only genes covered by at least 10 reads in a minimum of two replicates. Using a FDR adjusted p-value cut-off of 0.001, we retained as sex-biased the genes that had significant differences in expression between males and females in at least two of the three methods (Figure S3).

### Statistics

All statistical tests and figures were done using R (version 3.2.3, (Team 2016)).

## DATA ACCESS

Data are available in the NCBI database under the accession numbers SAMN12097880 to SAMN12097891

## ACKNOWLEDGMENTS

We thank the BGI for free sequencing thanks to their call for RNA-seq for medicinal plants. We thank Aline Muyle for advice with SEX-DETector and discussions. We thank Dr Tatyana Sukhorada, P.P. Lukyanenko Krasnodar Research and Development Institute of Agriculture for providing seeds of *C. sativa* cultivar ‘Zenitsa’. This project was supported through ANR grant ANR-14-CE19-0021-01 to G.A.B.M.

## Author Contributions

Conceptualization of the study: GABM, GK; Methodology: GABM, GK; Software: DP, BR, HB; Formal analysis: DP; Investigation: DP, OR, BR, HB, HH, JK, GK, GABM; Resources: OR, CF, GK; Data curation: CF; Writing - original draft: GABM, DP, JK; Writing – review and editing: all authors; Visualization: DP; Supervision: GABM, JK, GK; Project administration: GABM; Funding acquisition: GABM, GK.

## DISCLOSURE DECLARATION (including any conflicts of interest)

The authors declare no conflicts of interest

